# Strong selection is poorly aligned with genetic variation in *Ipomoea hederacea*: implications for divergence and constraint

**DOI:** 10.1101/2022.11.18.517124

**Authors:** Georgia Alexandria Henry, John R Stinchcombe

## Abstract

The multivariate evolution of populations is the result of the interactions between natural selection, drift, and the underlying genetic structure of the traits involved. Covariances among traits bias responses to selection, and the multivariate axis which describes the greatest genetic variation is expected to be aligned with patterns of divergence across populations. An exception to this expectation is when selection acts on trait combinations lacking genetic variance, which limits evolutionary change. Here we used a common garden field experiment of individuals from 57 populations of *Ipomoea hederacea* to characterize linear and nonlinear selection on five quantitative traits in the field. We then formally compare patterns of selection to previous estimates of within population genetic covariance structure (the G-matrix) and population divergence in these traits. We found that selection is poorly aligned with previous estimates of genetic covariance structure and population divergence. In addition, the trait combinations favoured by selection were generally lacking genetic variation, possessing approximately 15- 30% as much genetic variation as the most variable combination of traits. Our results suggest that patterns of population divergence are likely the result of the interplay between adaptive responses, correlated response, and selection favoring traits lacking genetic variation.

## Introduction

Understanding how populations change through time and space is a central goal of evolutionary biology. The strength and form of natural selection varies across landscapes and over time can lead to divergence among populations due to local adaptation (Endler 1977; Linhart and Grant 1996). While divergence among habitats or across gradients is often interpreted as adaptive (cf. Vasemägi 2006), natural selection acts in conjunction with other evolutionary forces such as genetic drift, which can contribute to divergence, and gene flow, which can act as a homogenizing force. The amount of genetic variation that exists within a population, and how that variation is organized among traits also influences how adaptive divergence proceeds (Lande and Arnold 1983; Falconer and Mackay 1996; Walsh and Blows 2009). The response to selection will depend on both natural selection acting on a population along with the variation within traits and the covariances among them (Antonovics 1976; Lande and Arnold 1983; Blows and Hoffmann 2005; Walsh and Blows 2009). Here we measure natural selection in the field on five quantitative traits and compare selection with estimates of clinal divergence and genetic covariances between traits, to formally evaluate the roles of selection and trait covariation in divergence.

The G-matrix (Lande 1979) summarizes the genetic variances and covariances among traits, with the genetic variances of traits along the diagonal of the matrix and covariances among those traits on the off-diagonals. Over short time scales **G** may bias response to selection of a population, thereby reducing the potential increase in fitness, if the direction of selection is not aligned with the major axes of variation in **G** (Schluter 1996; Walsh and Blows 2009). Additionally, the axis of greatest genetic variation, ***g****_max_*, is expected to be aligned with divergence among populations, as this axis biases the response to selection toward itself due to trait covariances (Schluter 1996), and via drift which also generates divergence along ***g****_max_* (Lande 1979; Arnold et al. 2001; Phillips et al. 2001). While absolute evolutionary constraints, whereby no evolutionary response to selection is possible due to the complete absence of genetic variation, are unlikely, any constraint due to the geometry of **G** will slow the evolutionary response to selection and, in cases where rapid evolutionary response is required such as during range expansion or in the face of climate change, populations may be at risk of extinction (Walsh and Blows 2009).

*Ipomoea hederacea* (ivyleaf morning glory), a weedy annual plant with an eastern range which stretches from approximately southern Pennsylvania and New Jersey through to Mexico, and has likely undergone evolutionarily recent, rapid range expansion (Campitelli and Stinchcombe 2014). The extent to which *Ipomoea hederacea*’s range in United States is due to a very recent invasion is contested, but herbaria records indicate it has persisted in its current range since at least the 1800’s (Bright-Emlen 1998). *Ipomoea hederacea* has genetic clines in both leaf shape, a Mendelian trait (Bright and Rausher 2008; Campitelli and Stinchcombe 2013a), and quantitative traits such as flowering time (Simonsen and Stinchcombe 2010; Stock et al. 2014). Stock and colleagues (2014) investigated latitudinal divergence among a suite of life history and morphological traits. They characterized the multivariate genetic divergence among populations and the multivariate genetic divergence with respect to latitude. Henry and Stinchcombe (2023) estimated G-matrices of four populations of *I. hederacea*, two from the northern-edge of the range and two from the core of the range, for the same suite of traits as Stock et al. (2014). Contrary to their expectations, they found no alignment between the population’s ***g****_maxes_* and the axis of latitudinal multivariate divergence (Cline_max_). These results suggested that strong selection has acted in a direction unaligned or orthogonal with **G**, leading to a lack of alignment between **G**, divergence, and selection. Alternatively, the observed lack of alignment may simply be due to unmeasured correlated traits having experienced selection, and the observed response being driven by the unmeasured traits (Lande and Arnold 1983).

To evaluate the relative roles of selection and potential correlated responses to selection in leading to divergence across *I. hederacea*’s range, we estimated selection in the field on a broad representation of populations. We grew *I. hederacea* plants from 57 populations from across its eastern North American distribution in the field, at the Koffler Scientific Reserve, King, Ontario. We estimated selection on five quantitative traits which capture aspects of growth and size, phenology, and reproductive biology. Specifically, we asked: 1) Are estimates of directional and non-linear selection correlated with previous estimates of **G** (Henry and Stinchcombe 2023) and divergence (Stock et al. 2014)? 2) How much genetic variance is there in the combination of traits under selection? And 3) Will patterns of trait variation and covariation reflected in **G** facilitate or constrain adaptation in northern populations of *I. hederacea*, thus potentially facilitating or constraining range expansion?

Our analyses necessarily combine data from different experimental environments: a field study (reported here) and two separate greenhouse common garden experiments (Stock et al. 2014, Henry and Stinchcombe 2022), which also differed in the number of populations and within-population samples. It is well known that the expression of phenotypic and genetic variation can vary by environment (e.g., Wood and Brodie 2015). Thus, we performed Krzanowski’s subspace analysis (Krzanowski 1979; following Aguirre et al. 2014) on the phenotypic variance-covariance matrices for northern and southern groups from the three experiments to evaluate the level of similarity between the phenotypic variation expressed across these experiments. We found overwhelming similarity among patterns of phenotypic covariation in all three experiments, strengthening our confidence that combining the results of these studies would yield a useful understanding of genetic variation, divergence, and selection in this system.

## Materials and Methods

### Study species and Natural History

*Ipomoea hederacea* (Convolvulaceae) is an annual vine found in disturbed habitats such as agricultural fields and roadsides in eastern USA, with a range that extends to southern Pennsylvania and mid-state New Jersey, USA. Plants germinate in early summer and grow and flower until a frost ends the growing season. Despite producing showy flowers, selfing rates are high (92-94%, Campitelli and Stinchcombe 2014).

### Quantitative Genetic and Field Experimental Design

We used replicate seeds, set by self-fertilization in a common greenhouse environment, of 343 maternal lines. Maternal lines were derived from 57 populations (1-10 lines per population, median = 6), gathered from 10 states in *I. hederacea’s* eastern North American distribution, spanning ∼7° of latitude (33.017681° to 40.340767° N). We specifically chose lines to increase overlap of previous studies of latitudinal divergence (Campitelli and Stinchcombe 2013; Stock et al. 2014), G-matrices within populations (Henry and Stinchcombe 2023), and measured the same phenotypes to maximize comparability across studies. We used 303 lines gathered by Campitelli and Stinchcombe (2013), from which we selected lines that had sufficient seeds and to maximize geographic localities. To this, we added 40 maternal lines, randomly chosen from Henry and Stinchcombe’s (2022) collections that had sufficient seeds. We note that Stock et al. (2014) also studied a subset of Campitelli and Stinchcombe (2013a) lines, and that our sample includes 19 of the 20 populations studied by Stock et al. (2014).

On 20 July 2021, we sowed 1024 scarified seeds from these maternal lines (339 lines with 3 seeds per line, 3 lines with 2 seeds per line, and 1 line with 1 seed) into 4” peat pots filled with Pro-Mix BX mycorrhizae soil in a glasshouse at ambient temperature and light. We transplanted pots into a recently ploughed old field three days later, with pots placed into the ground flush with field soil. We used a randomized block design of three spatial blocks in the old field (one replicate per line per block) and 1-meter grid spacing between individuals. We watered individuals thoroughly on the day they were transplanted into the ground but provided no further water supplementation. Approximately 90% of seeds germinated (917 of 1024), and plants were left unstaked and allowed to grow naturally among colonizing weeds, which included *Amaranthus palmeri*, several *Brassica spp.*, *Ambrosia artemisiifolia*, and *Capsella bursa-pastoris,* among others.

### Phenotypic Measurements

We measured a suite of size, phenology, and floral architecture traits that had been studied in prior investigations (Stock et al. 2014, Henry & Stinchcombe 2022): individual seed mass (g), early growth rate (leaves/day), days to onset of flowering (number of days after sowing), corolla width (mm) and anther-stigma distance (mm). We measured each individual’s seed mass prior to sowing. We calculated early growth rate as the difference in leaf number on two separate days divided by the number of intervening days. We performed our first leaf count survey when most plants had produced true leaves (22 days after sowing) and performed the second leaf count survey 12 days later. We conducted daily surveys to record the day on which each plant opened its first flower, which we used to measure corolla width and anther-stigma distance. We measured corolla width and anther-stigma distance using digital calipers (precision: ± 0.02 mm). We measured the distance from the lowest and highest anthers to the stigma and calculated the mean of the absolute distance to characterize anther-stigma distance.

We allowed individuals to grow, flower, set seed, and senesce naturally in the field. We collected the above-ground tissue four days after we observed widespread frost damage, when weather conditions were dry, on October 27th, 2021. Of the 1024 individuals sown at the beginning of the experiment, 650 had flowered by the end of the experiment, and 190 had set seeds. We counted the number of seeds produced from each individual, if any. We used the total seed set number as our proxy for fitness.

#### Subspace Analysis

One feature of our analyses is that they combine data from different experimental environments: a field study in a freshly ploughed field, and two separate greenhouse common garden experiments. Further, these studies differed in the number of populations sampled (4, 20, and 57), within-population replication of lines (50, 10, and ∼6), and total sample sizes (2137, 1467, and 650, for Henry & Stinchcombe 2022, Stock et al. 2014, and this study, respectively). To evaluate the similarity between the variation expressed across these experiments, we performed subspace analysis (Krzanowski 1979; following Aguirre et al. 2014) on the phenotypic variance-covariance matrices for northern and southern groups from the three experiments. We used > 38°N and < 36°N to characterize Northern and Southern populations, respectively (for 6 P-matrices total), as there were no populations sampled between 36-38°N. We used P-matrices because of the differences in quantitative genetic design and to maximize power. We then performed the subspace analysis for the P-matrices across all three experiments within each region and used the number of dimensions which explained at least 90% of the variation in the populations to construct the population subspaces, as per Aguirre et al (2014).

#### Analysis of Natural Selection

We standardized explanatory variables to have 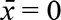 and σ = 1, which allowed us to compare the strength of selection acting on traits with different units (leaves/day, days, mm, etc.) and to compare the selection gradient directly with the G-matrices from Henry and Stinchcombe (2022) which were also estimated using standardized data. We divided the number of seeds set by each individual by the grand mean total seed set to calculate relative fitness; individuals which flowered, but did not set seed, had a relative fitness = 0, and were included in the analyses. Given the high selfing rates in this species (> 90%, Ennos 1981; Campitelli and Stinchcombe 2014), our use of seed set as a measure of fitness likely reflects both female and male fitness components, although any outcrossing in our experiment will reduce how much seed set reflects male fitness. To estimate selection on these traits we regressed relative fitness on our five traits using the *lme* function from the *nlme* package (Pinheiro et al. 2022) and included maternal line and field block as random effects to account for the non-independence of individuals of the same maternal line and environmental heterogeneity. We first estimated directional selection, ***β***, with all traits. To ensure that multicollinearity was not unduly affecting our estimates of selection (Mitchell-Olds and Shaw 1987; Chong et al. 2018), we calculated the variance inflation factor (VIF), which gives the amount of variance inflation in the model term due to multicollinearity with other model terms compared to a model without. While no clear boundaries exist for VIFs, values above 1 suggest some degree of correlation, and values above 5 are generally agreed to indicate a concerning level of multicollinearity (Menard 2002; James et al. 2021). We determined that no traits had a concerning degree of multicollinearity, as all traits had VIF values below 2 (Table S1). Thus, we retained all traits in the selection model. We also explored estimating selection on line means (Rausher 1992; Stinchcombe et al. 2002); we found very similar patterns of directional selection gradients, and as such present those in the Supplemental Material (Table S2) and focus on phenotypic estimates, while accounting for block and maternal line in the main text.

**Table 1:** Results from the linear mixed model of relative fitness (from total seed set) regressed on the five focal traits. Focal traits were variance-standardized. Family identity and field block are included as random effect terms.

| Trait | $\beta$ | Std. Error | df | t value | p-value |
| --- | --- | --- | --- | --- | --- |
| Seed mass | 0.2057 | 0.0946 | 341 | 2.1752 | 0.0303 |
| Growth rate | 0.4058 | 0.1078 | 341 | 3.766 | 0.0002 |
| Flowering time | -0.8583 | 0.1196 | 341 | -7.1735 | < 0.0001 |
| Corolla width | 0.2606 | 0.1139 | 341 | 2.2874 | 0.0228 |
| A-S distance | 0.0367 | 0.0995 | 341 | 0.3688 | 0.7125 |

**Table 2:**
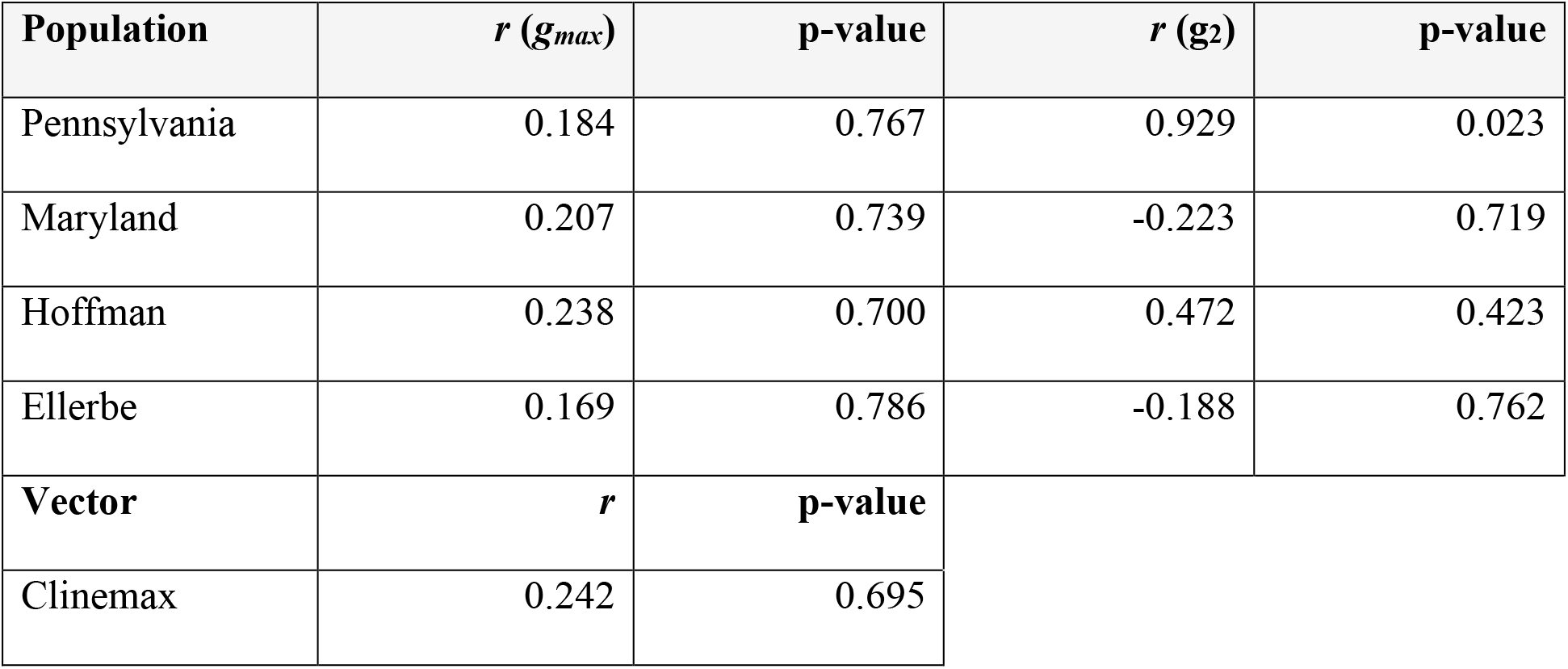
Pearson’s correlation coefficients and associated p-values for the estimated selection gradient and population ***g**_maxes_*. Correlation coefficient statistics are also given for the selection gradient and the axis of greatest multivariate divergence (*Cline_max_*).

To estimate nonlinear selection, we included quadratic terms and pairwise interactions of each trait. We multiplied the quadratic terms by a factor of 2, and constructed ***γ*** (Stinchcombe et al. 2008). To further aid in interpretation of the multivariate nonlinear selection gradient we performed canonical rotation of the axes (Phillips and Arnold 1989) and assessed significance of nonlinear selection via double-regression permutation testing (Reynolds et al. 2010), which incorporated sampling error in the null distribution. We then fit a thin plate spline using the *fields* package (Nychka et al. 2017) to the canonically rotated axes for visual interpretation.

#### Combining estimates of β, γ, G, and Divergence

We next used our field estimates of ***β*** to evaluate patterns of constraint and divergence, taking advantage of past studies using this species. First, we compared our field estimate of ***β*** to the axis of greatest latitudinal multivariate divergence described by Stock et al. (2014) to quantitatively test the hypothesis that there is a relationship between natural selection on these traits and their geographic divergence. Stock et al. (2014) measured the same phenotypes in a greenhouse common garden experiment; we used their original data to re-estimate latitudinal divergence on the 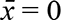 and σ = 1 scale. We used Pearson’s correlation coefficient to evaluate whether our estimated directional selection gradient and the axis of latitudinal multivariate divergence (*Cline_max_*) from their experiment were correlated with one another.

Second, we compared our estimates of ***β*** and ***γ*** to the G-matrices of four populations studied by Henry and Stinchcombe (2022), again in a common greenhouse environment. To do so, we examined the correlation between our estimated directional selection gradient and the axis of greatest variation (PC1, or ***g****_max_*) of each population and the correlation between ***β*** and the second eigenvector of the **G**s, *g_2_*, which describes the axis of second most variation. We also calculated the amount of genetic variation in the direction of selection, by projecting our estimates of ***β*** through each of the population G-matrices (***β****^T^***G*β***). To account for differences in size of the G-matrices, we standardized the projection by the eigenvalue of ***g****_max_*, such that a value of 1 would indicate that there is as much genetic variation in ***β***, as ***g****_max_*, the axis of greatest variation. As Henry and Stinchcombe (2022) used a Bayesian approach to estimate **G** we calculated all of the metrics described above for all posterior samples, and we report 95% Highest Posterior Density (HPD) intervals along with the mean values. We repeated this technique for the first two principal components of ***γ***, *m1* and *m2,* the axes of greatest nonlinear selection.

Third, we used our field estimates of ***β*** and ***γ*** to determine how much the genetic covariances described by Henry and Stinchcombe (2023) slow the rate of adaptation. We used the metrics described by Agrawal and Stinchcombe (2009), which compare the rate of adaptation with estimated genetic covariances to a scenario in which covariances are set to zero. We used both directional and nonlinear selection gradients (equation 2.3b in Agrawal and Stinchcombe 2009).

## Results

### Subspace analysis

The selection we estimate here was measured in a natural field environment, but the G-matrices (from Henry and Stinchcombe 2022) and the multivariate latitudinal axis of divergence (from Stock et al. 2014) were estimated in controlled glasshouse environments. We found that for both the northern and southern P-matrices the first two eigenvectors of the shared subspaces were completely shared among experiments and thus that they have a high degree of similarity (Table S7). While the environmental influences on each of the experimental groups could contribute to differences among them, the subspace analysis suggests that those differences are minimal with regards to phenotypic variation.

### Analysis of Natural Selection

We found overall strong directional selection, with all traits except anther-stigma distance exhibiting a significant relationship with relative fitness (Table 1). Selection favoured larger seed mass, faster growth rate, and increased corolla widths. Selection was acting against the number of days until the first flower, as has been described previously in this (Simonsen and Stinchcombe 2010,Campitelli and Stinchcombe 2013b) and other systems (Austen et al. 2017).

We found significant nonlinear selection acting on growth rate and flowering time with positive coefficients indicating the potential for disruptive selection (but see Mitchell-Olds and Shaw 1987), and negative correlational selection on seed mass and flowering time (Table S3). Because mixtures of disruptive and correlational selection can be difficult to interpret (Phillips and Arnold 1989; Simms 1990; Blows and Brooks 2003; McGoey and Stinchcombe 2009), we next performed a canonical rotation on the γ-matrix to evaluate the overall curvature of the nonlinear selection gradient (Phillips and Arnold 1989). Following Simms (1990) and Reynolds et al. (2010) we refit the trait data to the transformed axes and regressed fitness on the transformed trait values to obtain the approximate standard error of the eigenvalues of the rotated **γ**-matrix (Bisgaard and Ankenman 1996); we used permutation testing, following Reynolds et al. (2010) for significance testing, which evaluates the presence of overall nonlinear selection (Chenoweth et al. 2013).

**Table 3:**
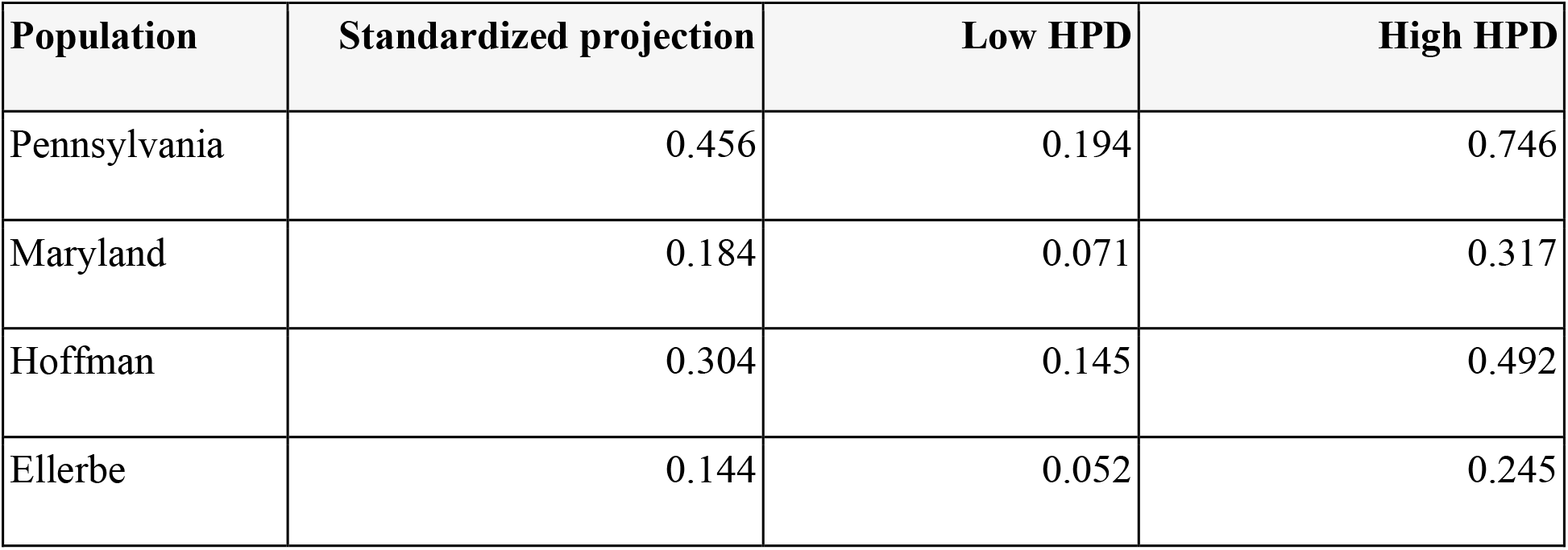
Projection of the direct selection gradient through each population **G**, standardized by the amount of variation in ***g****_max_*. 95% HPD intervals are given for each projection.

| Population | Standardized projection | Low HPD | High HPD |
| --- | --- | --- | --- |
| Pennsylvania | 0.456 | 0.194 | 0.746 |
| Maryland | 0.184 | 0.071 | 0.317 |
| Hoffman | 0.304 | 0.145 | 0.492 |
| Ellerbe | 0.144 | 0.052 | 0.245 |

We found that the first two canonical axes of **γ** are significant, with both signs being positive indicating a “bowl” shape (figure 1), with selection favouring negative values of both the first axis (*m1*) and second axis (*m2*). The dominant pattern along both axes was primarily linear (directional) selection with accelerating fitness benefits toward the lower values of *m1* and *m2*, rather than true disruptive selection with an intermediate fitness minimum (cf. Mitchell-Olds and Shaw 1987). Along *m1*, nonlinear selection favoured individuals with earlier flowering time (0.9780), larger corolla widths (−0.1442), larger initial seed mass (−0.1183), faster growth rate (− 0.0755), and smaller anther-stigma distances (0.0554) (Table S4). Along *m2* selection favoured individuals with faster growth rate (−0.9584), smaller anther-stigma distances (0.2109), larger corolla widths (−0.1399), later flowering onset (−0.1145), and larger initial seed mass (−0.0656). Together these axes suggest that while early flowering time is strongly associated with high fitness, selection also favours individuals with fast growth rates such that growth rate may be able to compensate for later flowering time.

**Figure 1:**
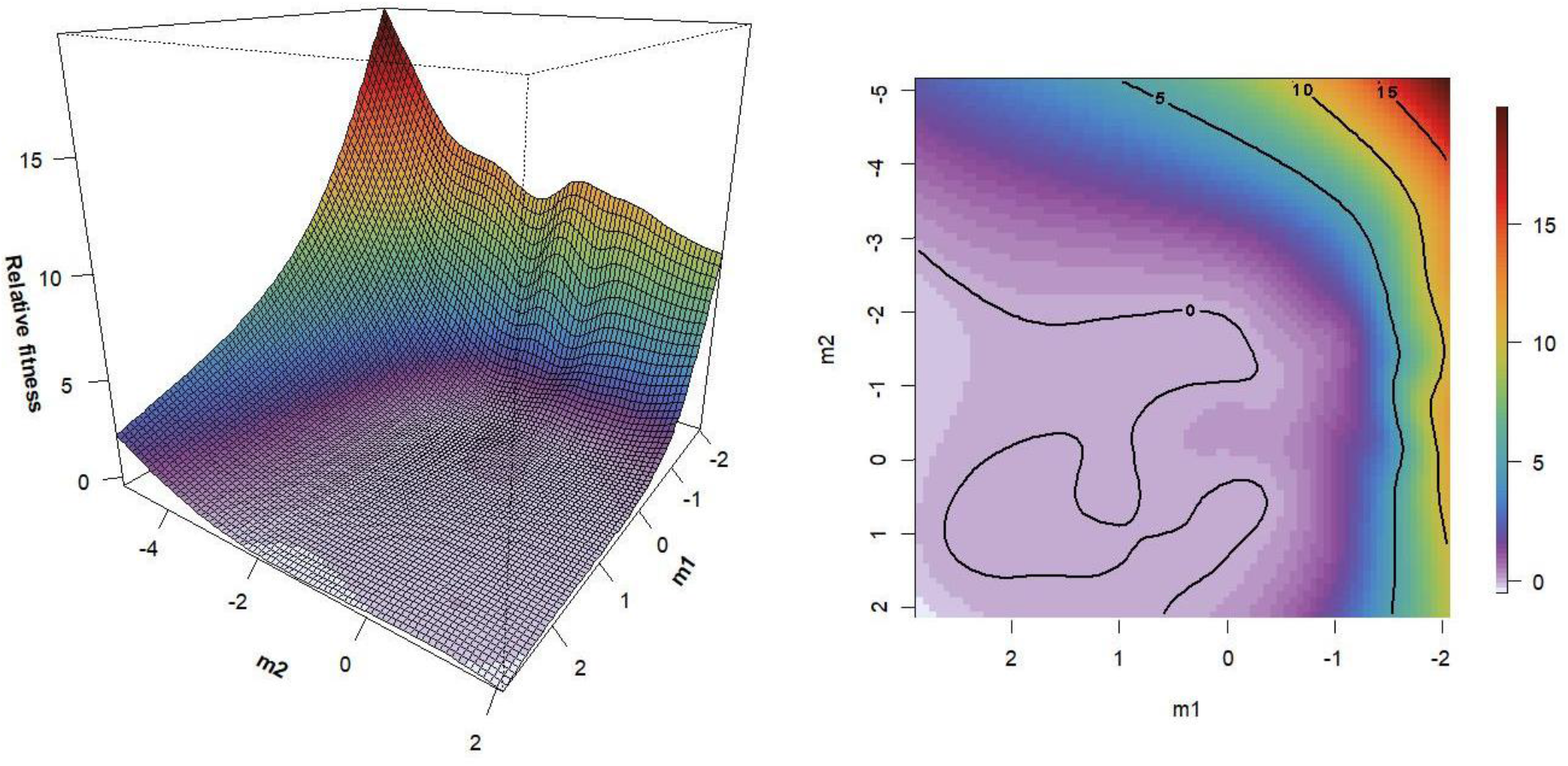
A) Perspective plot of a thin plate spline describing fitness with respect to the first two canonical axes of the γ-matrix. B) A contour plot of the same thin plate spline. High values of both axes have low fitness. Earlier flowering onset (*m1*) and faster early growth rate (*m2*) in combination is most strongly selected for, with regions of higher fitness along the extremes of both axes.

### Combining estimates of β, γ, G, and Divergence

The correlation between ***β*** and the axis of latitudinal divergence was low (*r* = 0.242) and not significantly different from the random expectation (p = 0.695) (Table 2). The axis of multivariate latitudinal divergence is thus not aligned with our northern selection gradient.

Our estimated selection gradient, ***β***, overall had low correlation with the populations ***g****_max_* and so was not significantly aligned to any population ***g****_ma_*_x_ (Table 2). Pennsylvania’s *g_2_* was significantly correlated with ***β*** (*r* = 0.93, p = 0.023), no other *g_2_* was significantly correlated with ***β*** (Table 2), although the correlation coefficient for ***β*** and Hoffman’s ***g****_2_* was moderate (*r* = 0.47). Expectedly, the amount of variation in the direction of selection is considerably reduced relative to ***g****_max_*. Pennsylvania, which had the greatest amount of variation in the direction of selection, had a ∼65% reduced variation when compared with ***g****_ma_*_x_, while Ellerbe, with the least variation, had an 85% reduction relative to ***g****_max_* (Table 3). We found comparably low levels of genetic variation present along *m1*, the first axis of the canonically rotated ***γ*** (Table S5). Although there is an increased level of genetic variation present along *m2* relative to that in the direction of ***β*** and *m1,* the reduction relative to ***g****_max_* is still considerable, within the range of 42-69% for all populations. Taken together, these results show that there is little genetic variance in the direction favoured by selection in our field study.

### The influence of **G** on adaptive evolution

We next evaluated how the trait covariances are predicted to influence the overall rate of adaptation. Of the four populations from Henry & Stinchcombe (2022) we found that three are predicted to be constrained overall by the organization of the population G-matrices when compared to a hypothetical with no covariances – Ellerbe by 56.36%, Hoffman by 18.22% and Maryland by 16.69% (Table S6). Pennsylvania, one of the northern populations, is predicted to have an evolutionary response slightly facilitated by the covariance structure of its traits, by 4.67%.

## Discussion

Using a broad sample of populations of *Ipomoea hederacea* from across a multivariate genetic cline in the eastern United States, we estimated directional and nonlinear selection in a natural setting on five quantitative traits. Through our comparisons of selection with G-matrices estimated from northern and southern populations along with the latitudinal axis of divergence, three major results emerged. First, the northern directional selection gradient, ***β***, is unaligned with the axis of among-population latitudinal divergence (Stock et al. 2014), and the G-matrices of two northern and two core populations estimated from a previous study (Henry & Stinchcombe 2022), as anticipated by Henry & Stinchcombe (2022). Second, the amount of genetic variation in **G** in the direction of ***β*** is quite low, again suggesting that selection favoured a combination of traits lacking variation. Third, the populations are likely constrained in their response to selection due to the structure of genetic variation in the populations, with one exception in the case of the Pennsylvania population, which we found has a slight facilitation in its expected evolutionary response. We discuss these results in the context of the relationship between ***β***, **G**, and divergence, and the role of **G** in facilitating or constraining adaptation to novel, northern environments, below.

### Selection, **G**, and divergence

**G** is expected to be aligned with divergence as it deflects the response to selection toward the axis of greatest variation (Schluter 1996). In *Ipomoea hederacea*, when previous estimates of **G** across four populations were compared with the axis of multivariate clinal divergence (Henry and Stinchcombe 2022) the predicted selection gradients were not aligned with the axis of greatest variation in **G** (***g****_max_*) for any population. Two possible explanations exist which could explain the discordance in the observed patterns: 1) that there was strong, directional selection acting in a direction largely orthogonal to **G**, and thus generating divergence unaligned with **G** or, 2) that there were traits missing from our estimates of **G** and divergence, such that if the missing traits had been included, greater alignment would have been detected (Henry and Stinchcombe 2022). Our results indicate that missing correlated traits need not be invoked to explain the absence of a relationship between divergence and ***g****_max_*. The lack of a relationship between ***g****_max_* and divergence is what would be predicted if selection were acting in directions lacking genetic variation, which is what we found. The northern selection gradient we estimated, as predicted in the first hypothesis, is not aligned with the majority of the variation in **G,** and there is little genetic variation in the direction of ***β***. While the possibility of missing traits can never be formally excluded – as it is always possible to measure more phenotypes, ad infinitum – we suggest the simplest explanation for the observed divergence is selection acting in conflict with the organization of genetic variation, and divergence occurring despite this due to the strength of selection. Given that directional and non-linear selection is acting on trait combinations with little but not absent genetic variation, the evolutionary response is expected to be slowed but not totally prevented (Blows and Hoffmann 2005; Blows and Walsh 2009).

While some studies have found among-population divergence occurring along the genetic line of least resistance (Hine et al. 2009; Costa E Silva et al. 2020; Royauté et al. 2020), our study adds to a body of literature demonstrating a lack of alignment at microevolutionary scales. Paccard et al. (2016) found *Arabidopsis lyrata* populations from across a latitudinal gradient had overall moderate to high angles between the axis of greatest multivariate divergence and population ***g****_maxes_*, demonstrating divergence without alignment with the genetic line of least resistance. Chenoweth and colleagues (2010) found consistent sexual selection in *Drosophila serrata* along a latitudinal cline, which acted orthogonally to their estimated population **G**s, and thus was constraining the response to sexual selection. The among-population divergence predicted was intermediate to ***β*** and **G**, indicating that in this instance, the reduction of variation along the axis of selection was leading to a bias in the response toward the genetic line of least resistance (Chenoweth et al. 2010). In a detailed analysis of the multivariate divergence among ecotypes and regions of a small sea snail, *Littorina saxatilis* throughout a well-studied hybrid zone, Garcia (2014) determined that divergence was aligned well with previous estimates of natural selection and the phenotypic variance-covariance matrix was not influential in the among-ecotype divergence. These studies, along with work presented here, highlight the need for a re-evaluation of the *a priori* expectation that among-population divergence should proceed along the axis of greatest genetic variation. While **G** will deflect responses to selection, the strength and orientation of selection are equally relevant. In contrast, over much longer time scales, adaptive divergence of rainbowfish species, *Melanotaenia spp*., in lake and stream environments appears to be unconstrained by the organization of **G** (McGuigan et al. 2005).

Despite most divergence among the two species likely being driven by drift, and aligned with ***g****_max_*, hydrodynamic differences which evolved were aligned with axes of limited genetic variation (McGuigan et al. 2005). In addition, as selection may be acting perpendicularly to **G**, the axis of among-population divergence may not serve as a good approximation of overall selection. For example, anther-stigma distance shows a stronger latitudinal cline than flowering time (Stock et al. 2014), yet our results (Table 1) show that flowering time is under much stronger selection; past studies from both this location (Simonsen and Stinchcombe 2010) and the center of the range (Campitelli and Stinchcombe 2013b) have also found similar patterns of strong selection favoring earlier flowering. We suggest exercising caution prior to making such simplifying assumptions without further interrogation of the system of interest.

### Constraint and facilitation beyond the range-edge

We measured natural selection beyond the species range edge (400 km north of the northernmost collections we have made), which gives a picture of how selection might act in the context of range expansion or facilitated migration. Given the observed stability in *I. hederacea*’s **G** across populations (Henry and Stinchcombe 2023) – which suggests it is stable enough over microevolutionary timescales to be useful in predictions of future evolutionary responses– we can use our estimates of **G** and ***β*** to project how the northern populations would evolve, along with the potential for range expansion. While the populations are not well aligned with selection, they have demonstrated divergence in mean trait values (Stock et al. 2014) indicating past responses to selection. Pennsylvania’s genetic (co)variation is somewhat aligned with selection, with the second axis of variation significantly correlated with ***β***. But overall, the G-matrices had reduced variation along the axis of directional selection, which will likely constrain responses to selection in Northern habitats. We did, however, find that the covariance structure of Pennsylvania may provide facilitation of the evolutionary response to selection using Agrawal and Stinchcombe’s extension of the multivariate breeder’s equation (Agrawal and Stinchcombe 2009). The other populations, including the other northern population (Maryland) were, in contrast, largely unaligned with ***β*** and evolutionarily constrained by its covariance structure, making it unclear how ubiquitous alignment with selection is among range-edge populations. While the orientation of **G** in the Pennsylvania population is fortuitous, whether alignment was in response to selection or whether it is simply due to drift is unknown.

The mixed results of the four populations emphasizes that whether a population may successfully establish outside of the current range will heavily depend on the source population. The mean population trait values along with the organization of **G** may both contribute to the persistence beyond the contemporary range limit (Blows and Walsh 2009). Populations near the current range-edge may not be source populations for any newly initiated populations outside the range, if long-distance or human-aided dispersal is common. (Campitelli and Stinchcombe 2014) interpreted population genetic patterns from sequencing data as consistent with long distance dispersal events, possibly mediated via agricultural equipment. Thus, the variability in alignment with northern selection gradients coupled with long-distance dispersal may be contributing to the maintenance of the current range limit.

## Conclusions

Overall, our study has demonstrated the benefit of combining both field studies, population sampling, and greenhouse quantitative genetic work in understanding the quantitative genetic architecture of wild populations. We found that selection is acting largely orthogonal to patterns of both within-population genetic variation, and divergence across the landscape. The among-population divergence observed in *Ipomoea hederacea*, rather than being purely adaptive, appear to be likely due to the interplay of poorly conditioned G-matrices (Henry and Stinchcombe 2022), selection favoring trait combinations with little genetic variance, and correlated responses to selection. The combination of strong natural selection unaligned with genetic variances-covariances may be more widely responsible for divergence than is currently appreciated. However, we have remarkably few case studies where landscape divergence, genetic covariance estimates, and estimates of natural selection are available for the same system– and fewer still for those with field estimates of selection. Obtaining these data for a diversity of systems will be a challenging, but important, ongoing empirical endeavor.

## References

Agrawal, A. F., and J. R. Stinchcombe. 2009. How much do genetic covariances alter the rate of adaptation? Proc. Biol. Sci. 276:1183–1191.

Aguirre, J. D., E. Hine, K. McGuigan, and M. W. Blows. 2014. Comparing G: multivariate analysis of genetic variation in multiple populations. Heredity 112:21–29.

Antonovics, J. 1976. The Nature of Limits to Natural Selection. Ann. Mo. Bot. Gard. 63:224–247. Missouri Botanical Garden Press.

Arnold, S. J., M. E. Pfrender, and A. G. Jones. 2001. The adaptive landscape as a conceptual bridge between micro-and macroevolution. Genetica 112–113:9–32.

Austen, E. J., L. Rowe, J. R. Stinchcombe, and J. R. K. Forrest. 2017. Explaining the apparent paradox of persistent selection for early flowering. New Phytol. 215:929–934.

Bisgaard, S., and B. Ankenman. 1996. Standard errors for the eigenvalues in second-order response surface models. Technometrics 38:238–246.

Blows, M., and B. Walsh. 2009. Spherical cows grazing in flatland: constraints to selection and adaptation. Pp. 83–101 in J. van der Werf, H.-U. Graser, R. Frankham, and C. Gondro, eds. Adaptation and Fitness in Animal Populations: Evolutionary and Breeding Perspectives on Genetic Resource Management. Springer.

Blows, M. W., and R. Brooks. 2003. Measuring nonlinear selection. Am. Nat. 162:815–820.

Blows, M. W., and A. A. Hoffmann. 2005. A reassessment of genetic limits to evolutionary change. Ecology 86:1371–1384.

Bright, K. L., and M. D. Rausher. 2008. Natural selection on a leaf-shape polymorphism in the Ivyleaf Morning Glory (*Ipomoea hederacea*). Evolution 62:1978–1990.

Campitelli, B. E., and J. R. Stinchcombe. 2013a. Natural selection maintains a single-locus leaf shape cline in Ivyleaf morning glory, *Ipomoea hederacea*. Mol. Ecol. 22:552–564.

Campitelli, B. E., and J. R. Stinchcombe. 2013b. Testing potential selective agents acting on leaf shape in *Ipomoea hederacea*: predictions based on an adaptive leaf shape cline. Ecol. Evol. 3:2409–2423.

Campitelli, B. E., and J. R. Stinchcombe. 2014. Population dynamics and evolutionary history of the weedy vine *Ipomoea hederacea* in North America. G3 4:1407–1416.

Chenoweth, S. F., J. Hunt, and H. D. Rundle. 2013. Analyzing and Comparing the Geometry of Individual Fitness Surfaces. in E. Svensson and R. Calsbeek, eds. The Adaptive Landscape in Evolutionary Biology. Oxford.

Chenoweth, S. F., H. D. Rundle, and M. W. Blows. 2010. The contribution of selection and genetic constraints to phenotypic divergence. Am. Nat. 175:186–196.

Chong, V. K., H. F. Fung, and J. R. Stinchcombe. 2018. A note on measuring natural selection on principal component scores. Evol Lett 2:272–280.

Costa E Silva, J., B.M. Potts, and P. A. Harrison. 2020. Population Divergence along a Genetic Line of Least Resistance in the Tree Species Eucalyptus globulus. Genes 11.

Endler, J. A. 1977. Geographic Variation, Speciation and Clines. Princeton University Press.

Ennos, R. A. 1981. Quantitative studies of the mating system in two sympatric species of Ipomoea (Convolvulaceae).

Falconer, D. S., and T. F. C. Mackay. 1996. Introduction to quantitative genetics. Fourth. Essex, UK: Longman Group.

Garcia, C. 2014. The divergence between ecotypes in a *Littorina saxatilis* hybrid zone is aligned with natural selection, not with intra-ecotype variation. Evol. Ecol. 28:793–810.

James, G., D. Witten, T. Hastie, and R. Tibshirani. 2021. Introduction to Statistical Learning: With Applications in R. Springer.

Henry, G. A., and J. R. Stinchcombe. 2023. G-matrix stability in clinally diverging populations of an annual weed. Evolution 77:49–62.

Hine, E., S. F. Chenoweth, H. D. Rundle, and M. W. Blows. 2009. Characterizing the evolution of genetic variance using genetic covariance tensors. Philos. Trans. R. Soc. Lond. B Biol. Sci. 364:1567–1578.

Krzanowski, W. J. 1979. Between-groups comparison of principal components. J. Am. Stat. Assoc. 74:703–707.

Lande, R. 1979. Quantitative genetic analysis of multivariate evolution, applied to brain: body size allometry. Evolution 33:402–416.

Lande, R., and S. J. Arnold. 1983. The measurement of selection on correlated characters. Evolution 37:1210–1226.

Linhart, Y. B., and M. C. Grant. 1996. Evolutionary significance of local genetic differentiation in plants. Annu. Rev. Ecol. Syst. 27:237–277. Ann. Revs.

McGoey, B. V., and J. R. Stinchcombe. 2009. Interspecific competition alters natural selection on shade avoidance phenotypes in *Impatiens capensis*. New Phytol. 183:880–891.

McGuigan, K., S. F. Chenoweth, and M. W. Blows. 2005. Phenotypic divergence along lines of genetic variance. Am. Nat. 165:32–43. Menard, S. 2002. Applied Logistic Regression Analysis. Sage.

Mitchell-Olds, T., and R. G. Shaw. 1987. Regression analysis of natural selection: statistical inference and biological interpretation. Evolution 41:1149–1161.

Nychka, D., R. Furrer, J. Paige, and S. Sain. 2017. fields: Tools for spatial data. R package version.

Phillips, P. C., and S. J. Arnold. 1989. Visualizing multivariate selection. Evolution 43:1209– 1222.

Phillips, P. C., M. C. Whitlock, and K. Fowler. 2001. Inbreeding changes the shape of the genetic covariance matrix in *Drosophila melanogaster*. Genetics 158:1137–1145.

Pinheiro J, Bates D, R Core Team. 2022. *nlme*: Linear and nonlinear mixed effects models. R package version.

Rausher, M. D. 1992. The measurement of selection on quantitative traits: biases due to environmental covariances between traits and fitness. Evolution 46:616–626.

Reynolds, R. J., D. K. Childers, and N. M. Pajewski. 2010. The distribution and hypothesis testing of eigenvalues from the canonical analysis of the gamma matrix of quadratic and correlational selection gradients. Evolution 64:1076–1085.

Royauté, R., A. Hedrick, and N. A. Dochtermann. 2020. Behavioural syndromes shape evolutionary trajectories via conserved genetic architecture. Proc. Biol. Sci. 287:20200183.

Schluter, D. 1996. Adaptive radiation along genetic lines of least resistance. Evolution 50:1766– 1774.

Simms, E. L. 1990. Examining selection on the multivariate phenotype: plant resistance to herbivores. Evolution 44:1177–1188.

Simonsen, A. K., and J. R. Stinchcombe. 2010. Quantifying Evolutionary Genetic Constraints in the Ivyleaf Morning Glory, *Ipomoea hederacea*. Int. J. Plant Sci. 171:972–986.

Stinchcombe, J. R., A. F. Agrawal, P. A. Hohenlohe, S. J. Arnold, and M. W. Blows. 2008. Estimating nonlinear selection gradients using quadratic regression coefficients: double or nothing? Evolution 62:2435–2440.

Stinchcombe, J. R., M. T. Rutter, D. S. Burdick, P. Tiffin, M. D. Rausher, and R. Mauricio. 2002. Testing for environmentally induced bias in phenotypic estimates of natural selection: theory and practice. Am. Nat. 160:511–523.

Stock, A. J., B. E. Campitelli, and J. R. Stinchcombe. 2014. Quantitative genetic variance and multivariate clines in the Ivyleaf morning glory, *Ipomoea hederacea*. Philos. Trans. R. Soc. Lond. B Biol. Sci. 369:20130259.

Vasemägi, A. 2006. The adaptive hypothesis of clinal variation revisited: single-locus clines as a result of spatially restricted gene flow. Genetics 173:2411–2414.

Walsh, B., and M. W. Blows. 2009. Abundant genetic variation + strong selection = multivariate genetic constraints: a geometric view of adaptation. Annu. Rev. Ecol. Evol. Syst. 40:41–59.

Wood, C. W., and E. D. Brodie 3rd. 2015. Environmental effects on the structure of the G- matrix. Evolution 69:2927–2940.

