## Supplementary material for "Strong selection is poorly aligned with genetic variation in *Ipomoea hederacea*: implications for divergence and constraint"

### Supplementary Information

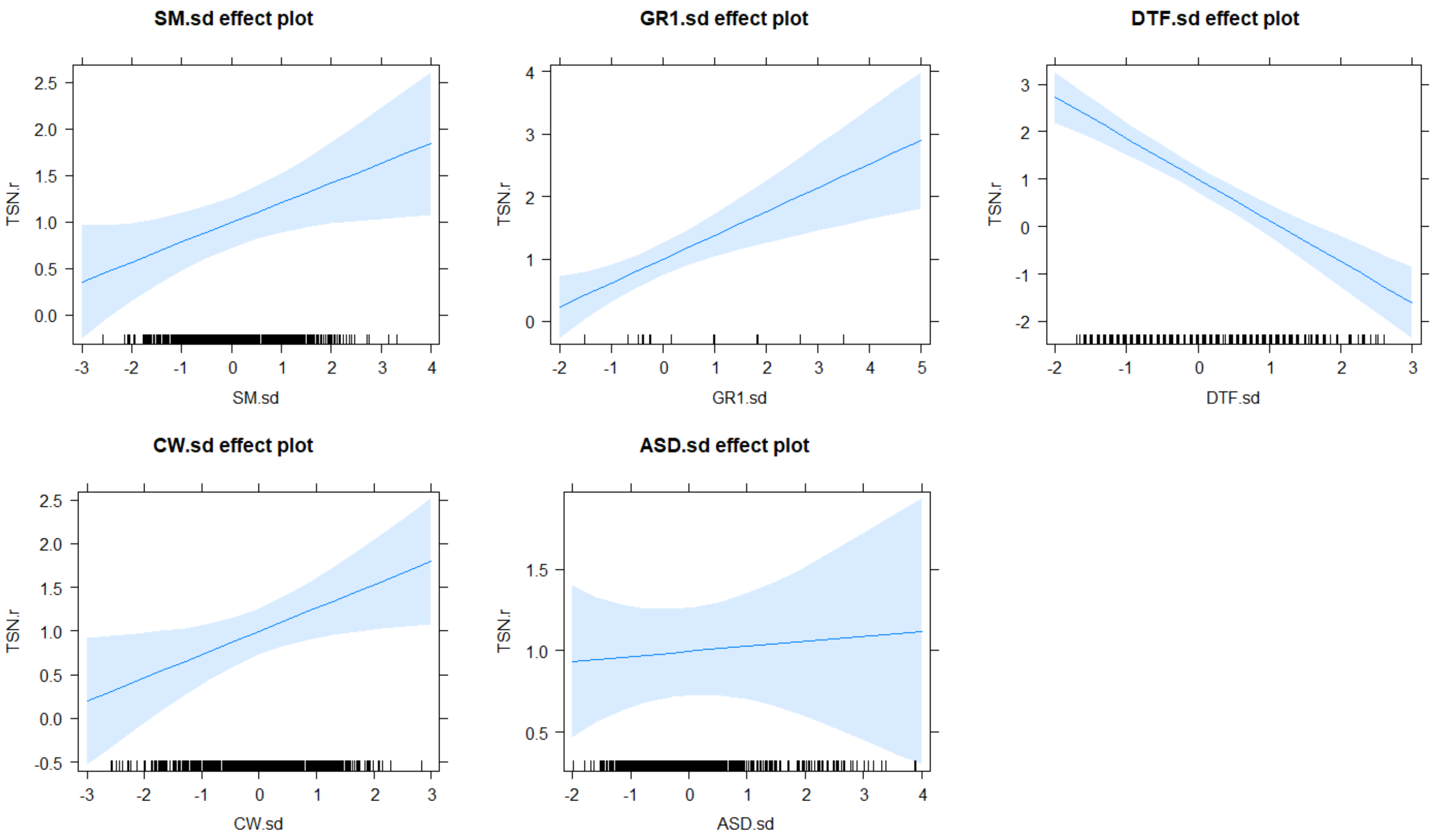


**Figure S1**: partial effect plots for each trait on fitness (relative total seed number) conditioned on the remaining variables in the model.

**Table S1:** Variance inflation factors for the directional selection model.

| Seed Mass | Growth Rate | Flowering Time | Corolla Width | A-S Distance |
|---:|---:|---:|---:|---:|
| 1.032512 | 1.380356 | 1.674856 | 1.533698 | 1.177777 |

**Table S2:** Directional selection estimate using line means.

| Trait | $\beta$ | Std. Error | *df* | F statistic | p-value |
|---|---:|---:|---:|---:|---:|
| Seed mass | 0.218 | 0.099 | 298 | 2.208 | 0.0280 |
| Growth rate | 0.159 | 0.106 | 298 | 1.499 | 0.1349 |
| Flowering time | -0.723 | 0.117 | 298 | -6.179 | < 0.0001 |
| Corolla width | 0.346 | 0.115 | 298 | 3.007 | 0.0029 |
| A-S distance | -0.053 | 0.104 | 298 | -0.512 | 0.6091 |

**Table S3:** M-matrix, the canonically rotated γ-matrix.

| | *m1* | *m2* | *m3* | *m4* | *m5* |
|---|---|---|---|---|---|
| Seed mass | -0.118 | -0.066 | -0.168 | 0.942 | 0.259 |
| Growth rate | -0.075 | -0.958 | 0.105 | 0.012 | -0.254 |
| Flowering time | 0.978 | -0.114 | 0.024 | 0.077 | 0.155 |
| Corolla width | -0.144 | -0.140 | 0.555 | -0.147 | 0.793 |
| A-S distance | 0.055 | 0.211 | 0.807 | 0.293 | -0.464 |
| Eigenvalue | 1.626 | 0.712 | 0.050 | -0.128 | -0.300 |

**Table S4:** Projection of first two principal components of **M** on population G-matrices standardized by the eigenvalue of $g_{max}$. Upper and lower bounds of the 95% HPD intervals (lower and upper CI) also shown. Overall the amount of variation along *m1* is comparable to that of $\beta$. In all populations there is similar or greater variation along *m2* than $\beta$, although considerably less than the maximum.

| Population | *m1* | lower CI | upper CI | *m2* | lower CI | upper CI |
|---|---|---|---|---|---|---|
| Pennsylvania | 0.393 | 0.185 | 0.676 | 0.383 | 0.13 | 0.65 |
| Maryland | 0.104 | 0.044 | 0.167 | 0.473 | 0.183 | 0.783 |
| Hoffman | 0.251 | 0.116 | 0.414 | 0.311 | 0.139 | 0.518 |
| Ellerbe | 0.155 | 0.073 | 0.242 | 0.577 | 0.302 | 0.865 |

**Table S5:** Krzanowski's lambdas from subspace analyses of P-matrices calculated from Stock et al. (2014), Henry and Stinchcombe (2022) and this paper. Due to differences in experimental design the populations were split into northern and southern groups and subspace analysis of the experimental P-matrices was conducted within those regions. In both cases the first two axes of the shared subspace were completely shared among experiments.

| Region | *h1* | *h2* | *h3* | *h4* |
|---|---|---|---|---|
| North | 3.000 | 3.000 | 2.447 | 1.680 |
| South | 3.000 | 3.000 | 2.779 | 0.992 |
